# A novel antibody against the furin cleavage site of SARS-CoV-2 spike protein: effects on proteolytic cleavage and ACE2 binding

**DOI:** 10.1101/2021.02.09.430451

**Authors:** Michael G. Spelios, Jeanne M. Capanelli, Adam W. Li

## Abstract

SARS-CoV-2 harbors a unique S1/S2 furin cleavage site within its spike protein, which can be cleaved by furin and other proprotein convertases. Proteolytic activation of SARS-CoV-2 spike protein at the S1/S2 boundary facilitates interaction with host ACE2 receptor for cell entry. To address this, high titer antibody was generated against the SARS-CoV-2-specific furin motif. Using a series of innovative ELISA-based assays, this furin site blocking antibody displayed high sensitivity and specificity for the S1/S2 furin cleavage site, and demonstrated effective blockage of both enzyme-mediated cleavage and spike-ACE2 interaction. The results suggest that immunological blocking of the furin cleavage site may afford a suitable approach to stem proteolytic activation of SARS-CoV-2 spike protein and curtail viral infectivity.

## Introduction

Severe acute respiratory syndrome coronavirus 2 (SARS-CoV-2) was first recognized in the beginning of 2020 and is responsible for the present COVID-19 pandemic. SARS-CoV-2 consists of a positive-sense single-stranded RNA genome and 4 different types of structural proteins.

The N, or nucleocapsid, protein encapsidates the genome, while the S (spike), E (envelope), and M (membrane) proteins comprise the surrounding lipid bilayer envelope. Of particular appeal is the S protein, which enables viral infection via angiotensin-converting enzyme 2 (ACE2) receptor recognition and membrane fusion, making this structural protein an ideal target for therapeutic intervention.

The S protein is composed of two subunits, S1 and S2. Within the S1 subunit is a receptor-binding domain (RBD) that recognizes and binds to the ACE2 receptor. The S protein also harbors a furin cleavage site at the boundary between the subunits. The complete SARS-CoV-2 furin cleavage site has been characterized as a 20 amino acid motif corresponding to the amino acid sequence ^A^672-^S^691 of SARS-CoV-2 spike protein (Figure 1), with one core region SPRRAR│SV (8 amino acids, ^S^680-^V^687) and two flanking solvent-accessible regions (8 amino acids, ^A^672–^N^679, and 4 amino acids, ^A^688-^S^691). The core region is very unique as its ^R^683 and ^A^684 positions are positively-charged (Arg) and hydrophobic (Ala) residues, respectively, which could be cleaved by the proprotein convertase (PC) furin and/or furin-like PCs secreted from host cells and bacteria in the airway epithelium. Furin and furin-like PCs, such as PC5/6A and PACE4, are proven to be cleavage region sequence-specific, and these PCs exhibit widespread tissue distribution. With this unique furin cleavage site, such distribution may explain why COVID-19 causes damage in multiple organs. Thus, the importance of blocking SARS-CoV-2 S1/S2 site cleavage caused by furin or facilitating protease activity is emphasized by the fact that cleavage of the S protein at the S1/S2 site has been documented as essential for SARS-CoV-2 binding to the host ACE2 receptor, cell-cell fusion, and infection of human lung cells [**1**,**2**].

**Figure 1.**
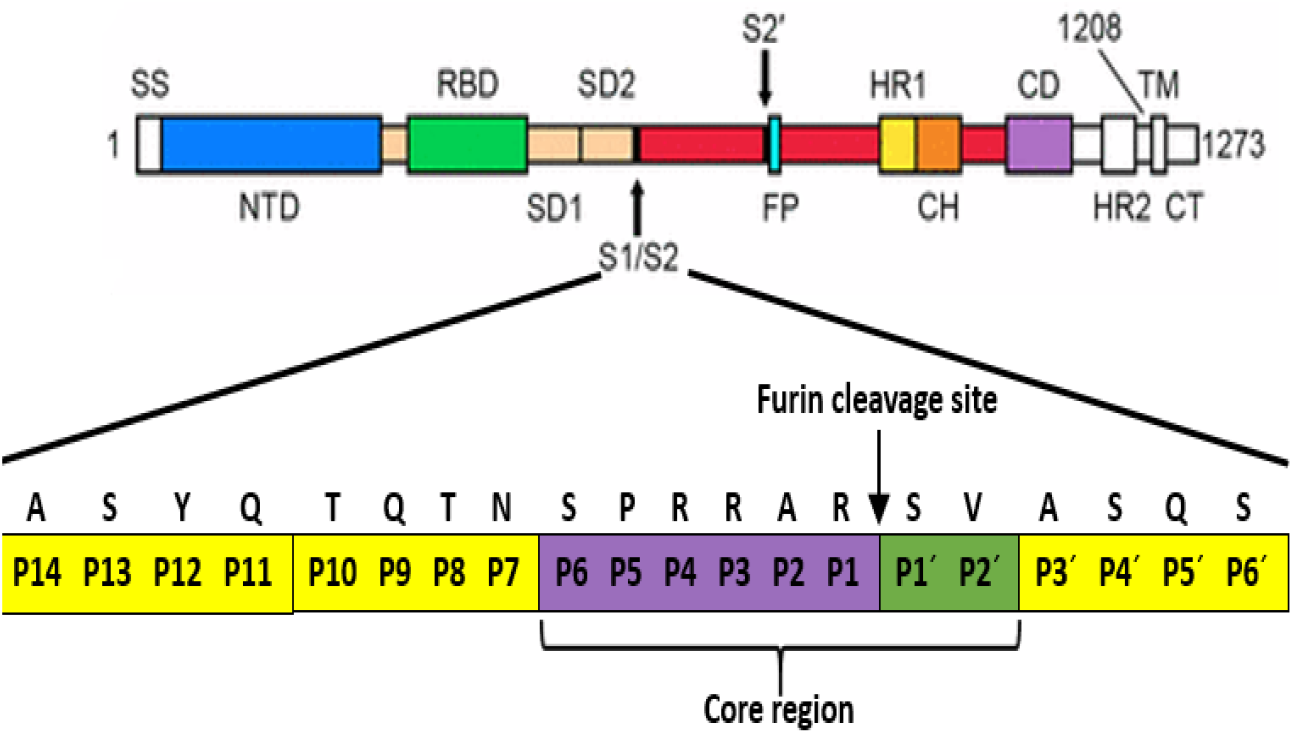
Structure of the SARS-CoV-2 spike protein, including the location of the furin cleavage site at the boundary between the S1 and S2 subunits.

Due to the correlation between PC-mediated cleavage and S protein activation, we hypothesized that blockage of the furin cleavage site could potentially subvert SARS-CoV-2 infection by hindering enzymatic action and S protein-ACE2 interaction. To address this, we generated a novel antibody with high sensitivity and specificity for the furin cleavage site of SARS-CoV-2 spike protein. The antibody was then employed in a series of innovative in vitro assays where it was observed to efficiently block a) cleavage by purified enzymes and human specimens, and b) binding to ACE2. To our knowledge, this is the first time that such an antibody-based tactic directed against the target of SARS-CoV-2-specific PC activity has been established.

## Methods

### Generation of furin site blocking antibody (fbAB)

Antigen consisted of the SARS-CoV-2-specific furin motif (20 aa; Figure 1) conjugated to keyhole limpet hemocyanin (KLH). The peptide-KLH conjugate (0.5 mg) was injected into a rabbit, followed by a boost (3 × 0.25 mg) over 2 weeks. Serum was collected and antibody was purified with protein A columns.

### Quantification of fbAB titer

To test the titer of the generated fbAB, the antigen was coated onto high protein binding, polystyrene, 8-well microplate strips at a concentration of 200 ng/well with 0.1 M NaCO_3_. The strips were incubated for 2 h at 37°C for coating and then blocked with 2% BSA for 1 h at 37°C. After washing the strips for 3 times with PBS-T, fbAB was added into the wells at the indicated dilutions (prepared with PBS-T) and incubated for 1 h at RT. After washing for 4 times, anti-rabbit IgG-HRP (EpiGentek) (50 µl, 1:2000) was added and incubated for 30 min at RT. After washing for 4 times, 100 µl of TMB solution (EMD Millipore Corp.) were added per well and blue color development was monitored for 2-10 min. The reaction was stopped with an equal volume of 1 M HCl and the optical density was measured with a microplate reader (MRX-TC Revelation, Dynex Technologies) at a wavelength of 450 nm.

### Sensitivity of fbAB recognition

A SARS-CoV-2 protein (Sino Biological), containing the S1/S2 boundary furin site and tagged with polyhistidine (His) at the N-terminal, was added at different concentrations (0.01 - 100 ng/well) into nickel-nitrilotriacetic acid (Ni-NTA)-coated polystyrene, 8-well microplate strips (Fisher) and incubated for 45 min at RT. After washing the strips for 2 times with PBS-T, fbAB (50 µl, 1:2000) was added and incubated for 1 h at RT. After washing for 3 times, anti-rabbit IgG-HRP (50 µl, 1:2000) was added and incubated for 30 min at RT. After washing for 4 times, 100 µl of TMB solution were added per well and blue color development was monitored for 2-10 min. The reaction was stopped with an equal volume of 1 M HCl and the optical density was measured with a microplate reader at a wavelength of 450 nm.

### Specificity of fbAB recognition

His-tagged SARS-CoV-2 protein containing the S1/S2 boundary furin site, His-tagged peptide containing the SARS-CoV-2-specific furin motif (EpiGentek), and His-tagged SARS-CoV-2 S1 RBD protein lacking the S1/S2 boundary furin site (EpiGentek) were added at a concentration of 10 ng/well to the Ni-NTA-coated strips and incubated for 1h at 37°C. After washing the strips for 2 times with PBS-T, fbAB (50 µl) was added at 1:1000 or 1:5000 dilutions and incubated for 1 h at RT. After washing for 3 times, anti-rabbit IgG-HRP (50 µl, 1:2000) was added and incubated for 30 min at RT. After washing for 4 times, 100 µl of TMB solution were added per well and blue color development was monitored for 2-10 min. The reaction was stopped with an equal volume of 1 M HCl and the optical density was measured with a microplate reader at a wavelength of 450 nm.

### Immunoprecipitation by fbAB

fbAB was coated onto the 8-well microplate strips at a concentration of 200 ng/well with 0.1 M NaCO_3_. The strips were incubated for 2 h at 37°C for coating, washed for 3 times with PBS-T, and then blocked with 2% BSA for 1 h at 37°C. After washing the strips for 3 times, His-tagged SARS-CoV-2 protein containing the S1/S2 boundary furin site or His-tagged peptide containing the SARS-CoV-2-specific furin motif were added at different concentrations to the fbAB-coated wells and incubated for 2 h at RT. After washing for 3 times, Ni-NTA-HRP (Millipore) (50 µl, 1:4000) was added and incubated for 30 min at RT. After washing for 4 times, 100 µl of TMB solution were added per well and blue color development was monitored for 2-10 min. The reaction was stopped with an equal volume of 1 M HCl and the optical density was measured with a microplate reader at a wavelength of 450 nm.

### fbAB blockage of furin-mediated cleavage

A peptide (EpiGentek) containing the SARS-CoV-2-specific furin motif and tagged with His and biotin at the N- and C-terminals, respectively, was added at a concentration of 10 ng/well to the Ni-NTA-coated strips and incubated for 1 h at RT. After washing the strips for 3 times with PBS-T, fbAB was added at a concentration of 200 ng/well and incubated for 1 h at 37°C. A SARS-CoV-2 neutralization antibody (EpiGentek), which targets the spike RBD, was used as a control. After washing for 3 times, purified proprotein convertase furin (New England Biolabs) was added at different concentrations (2-4 U/well) and incubated for 25 min at 37°C. Protease cleavage (PC) assay buffer (EpiGentek) was used to prepare the furin solutions. After washing for 4 times, streptavidin-HRP (100 µl, 1:5000) was added and incubated for 15 min at RT. After washing for 4 times, 100 µl of TMB solution were added per well and blue color development was monitored for 2-10 min. The reaction was stopped with an equal volume of 1 M HCl and the optical density was measured with a microplate reader at a wavelength of 450 nm.

### fbAB blockage of trypsin-mediated cleavage

His- and biotin-tagged peptide containing the SARS-CoV-2-specific furin motif was added at a concentration of 10 ng/well to the Ni-NTA-coated wells of the 8-well microplate strips and incubated for 1 h at RT. After washing the strips for 3 times with PBS-T, fbAB was added at a concentration of 200 ng/well and incubated for 1 h at 37°C. After washing for 3 times, serine protease trypsin (Sigma) was added at different concentrations (10-20 ng/well, prepared with PC assay buffer) and incubated for 25 min at 37°C. After washing for 4 times, streptavidin-HRP (100 µl, 1:5000) was added and incubated for 15 min at RT. After washing for 4 times, 100 µl of TMB solution were added per well and blue color development was monitored for 2-10 min. The reaction was stopped with an equal volume of 1 M HCl and the optical density was measured with a microplate reader at a wavelength of 450 nm.

### fbAB blockage of human nasal swab-mediated cleavage

Collection of nasal swab samples from healthy uninfected volunteers was in accordance with the standard CDC nasal swab collection protocol. The collected samples were released into 300 µl of PC assay buffer by rotating the swab in the buffer for 30 sec.

His- and biotin-tagged peptide containing the SARS-CoV-2-specific furin motif was added at a concentration of 10 ng/well to the Ni-NTA-coated wells of the 8-well microplate strips and incubated for 1 h at RT. After washing the strips for 3 times with PBS-T, fbAB was added at a concentration of 200 ng/well and incubated for 1 h at 37°C. After washing for 3 times, 20-30 µl of nasal swab sample solution were added per well and incubated for 25 min at 37°C. After washing for 4 times, streptavidin-HRP (100 µl, 1:5000) was added and incubated for 15 min at RT. After washing for 4 times, 100 µl of TMB solution were added per well and blue color development was monitored for 2-10 min. The reaction was stopped with an equal volume of 1 M HCl and the optical density was measured with a microplate reader at a wavelength of 450 nm.

### fbAB blockage of spike-ACE2 binding

Untagged SARS-CoV-2 spike protein (GenScript) containing the S1/S2 boundary furin site was coated onto the high protein binding, polystyrene, 8-well microplate strips at a concentration of 50 ng/well with 0.1 M NaCO_3_. The strips were incubated for 2 h at 37°C for coating and then blocked with 2% BSA for 1 h at 37°C. After washing the strips for 3 times with PBS-T, fbAB was added into the wells at the indicated concentrations and incubated for 1 h at 37°C. After washing for 3 times, purified His-tagged ACE2 (EpiGentek) was added at a concentration of 100 ng/well (prepared with PBS) and incubated for 1 h at 37°C. After washing for 3 times, Ni-NTA-HRP (50 µl, 1:4000) was added and incubated for 30 min RT. After washing for 4 times, 100 µl of TMB solution were added per well and blue color development was monitored for 2-10 min. The reaction was stopped with an equal volume of 1 M HCl and the optical density was measured with a microplate reader at a wavelength of 450 nm.

## Results

After generation of furin site blocking antibody (fbAB) raised against an antigen consisting of the 20 amino acid SARS-CoV-2-specific furin motif (Figure 1), the fbAB titer was evaluated using an ELISA-based colorimetric detection system with antigen-coated microplates. As shown in Figure 2, when compared with normal serum, fbAB purified from serum of antigen-injected host displayed a strong signal intensity (OD >2.5) at 8000X dilution. The optical density decreased linearly over successive 2-fold serial dilutions, but a high signal intensity (OD >1) was still observed for fbAB diluted as much as 128000X. The results indicate that high amounts of fbAB against the furin motif of SARS-CoV-2 spike protein could be generated from antigen-injected host and subsequently detected with this novel colorimetric assay.

**Figure 2.**
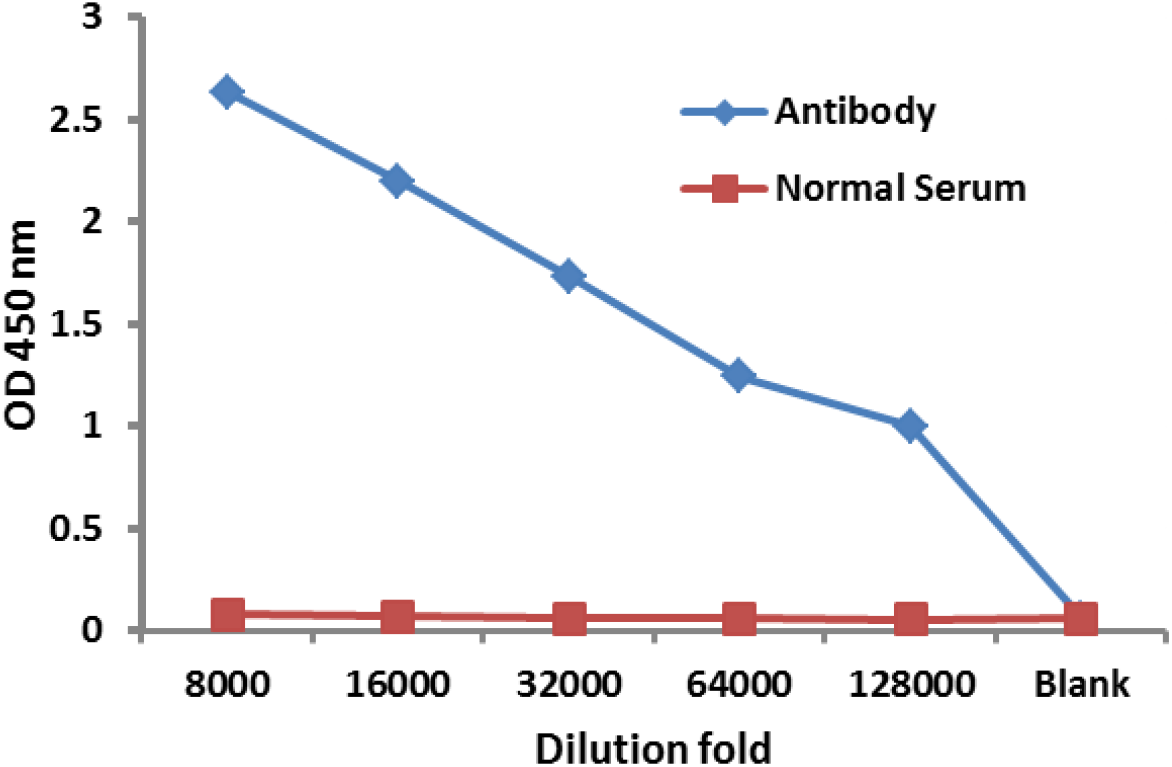
fbAB titer of recognizing the antigen containing the SARS-CoV-2-specific furin motif. Antigen concentration = 200 ng/well.

The sensitivity of fbAB recognizing the S1/S2 boundary furin site of SARS-CoV-2 spike protein was determined by incubating fbAB in nickel-coated microplate wells bound with different amounts of His-tagged target protein. Capture of the well-bound target by fbAB was then measured colorimetrically using ELISA-based detection. As shown in Figure 3, the target protein displayed a nice dose-response, with signal intensity increasing linearly up to a concentration of 100 ng/well. As low as 0.01 ng/well of the protein was detected with this assay, indicating high sensitivity of fbAB for SARS-CoV-2-specific furin motif recognition.

**Figure 3.**
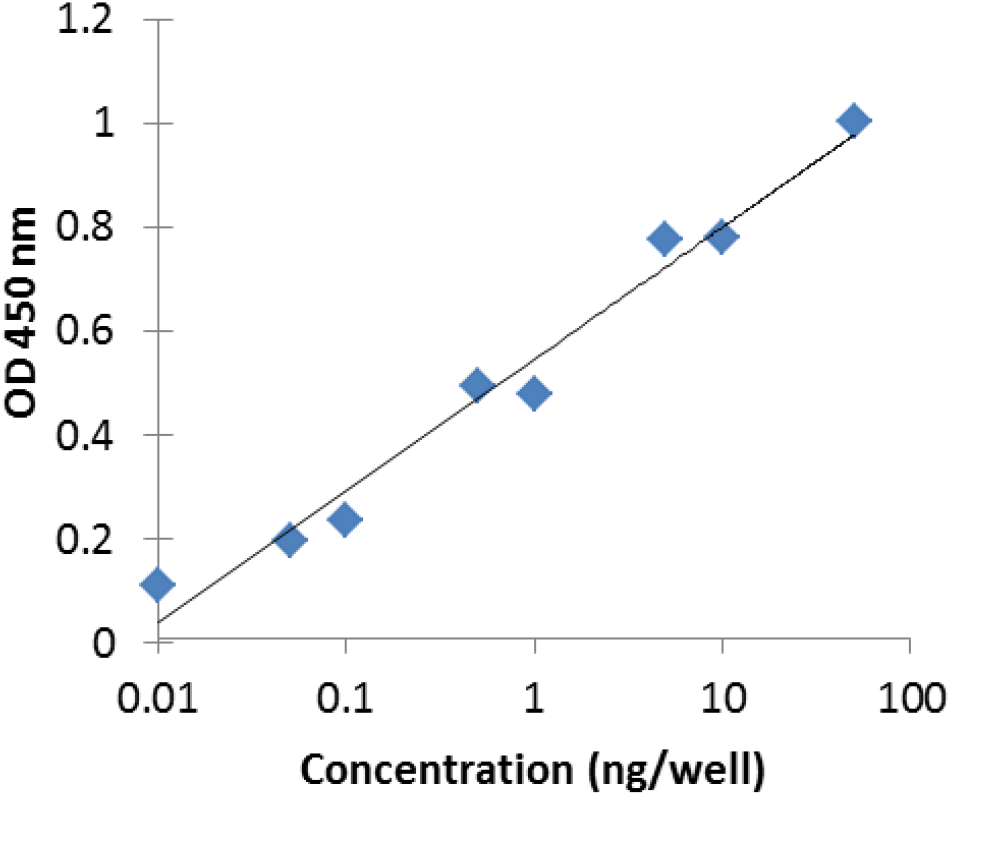
Sensitivity of fbAB recognizing the antigen containing the SARS-CoV-2-specific furin motif.

The specificity of fbAB-spike interaction was assessed by incubating antibody in wells coated with either: SARS-CoV-2 spike protein containing the S1/S2 boundary furin site; a peptide containing the SARS-CoV-2-specific furin motif; or SARS-CoV-2 S1 RBD protein. As shown in Figure 4, fbAB displayed a strong binding interaction with the furin cleavage site of both spike protein and peptide. The interaction was highly specific as the SARS-CoV-2 S1 RBD protein, which lacks the S1/S2 boundary furin site, failed to elicit a detectable signal. The S protein and peptide were further used to assess the immunoprecipitation efficiency of fbAB in antibody-coated wells. Figure 5 shows the efficient immunocapture of both S protein and peptide by fbAB in a concentration-dependent manner. The results indicate that synthesized peptide is the same as biological S protein for use as an assay substrate, as both the peptide and full-length S protein were strongly bound by fbAB. Thus, synthesized peptide was utilized in the follow-up cleavage blockage assays.

**Figure 4.**
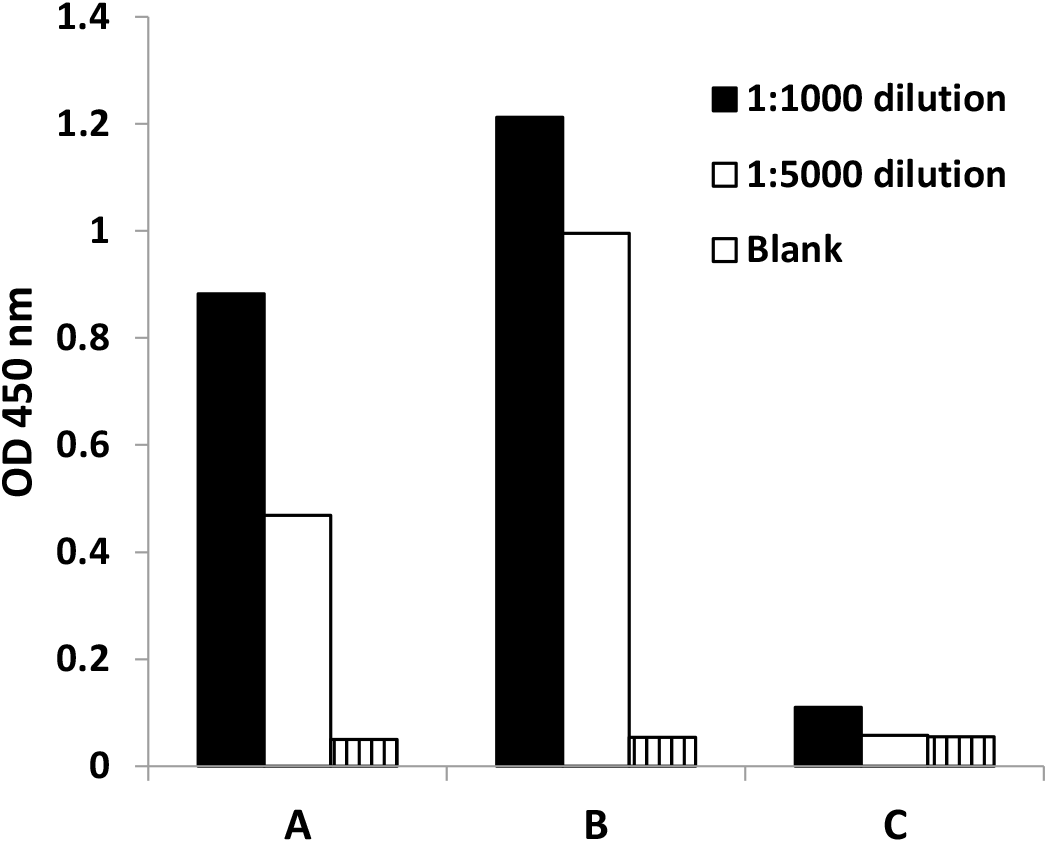
Specificity of fbAB recognizing the SARS-CoV-2 spike protein containing the S1/S2 boundary furin site or a peptide containing the SARS-CoV-2-specific furin motif. A: SARS-CoV-2 protein containing the S1/S2 boundary furin site; B: Peptide containing the SARS-CoV-2-specific furin motif; C: SARS-CoV-2 S1 RBD protein lacking the S1/S2 boundary furin site.

**Figure 5.**
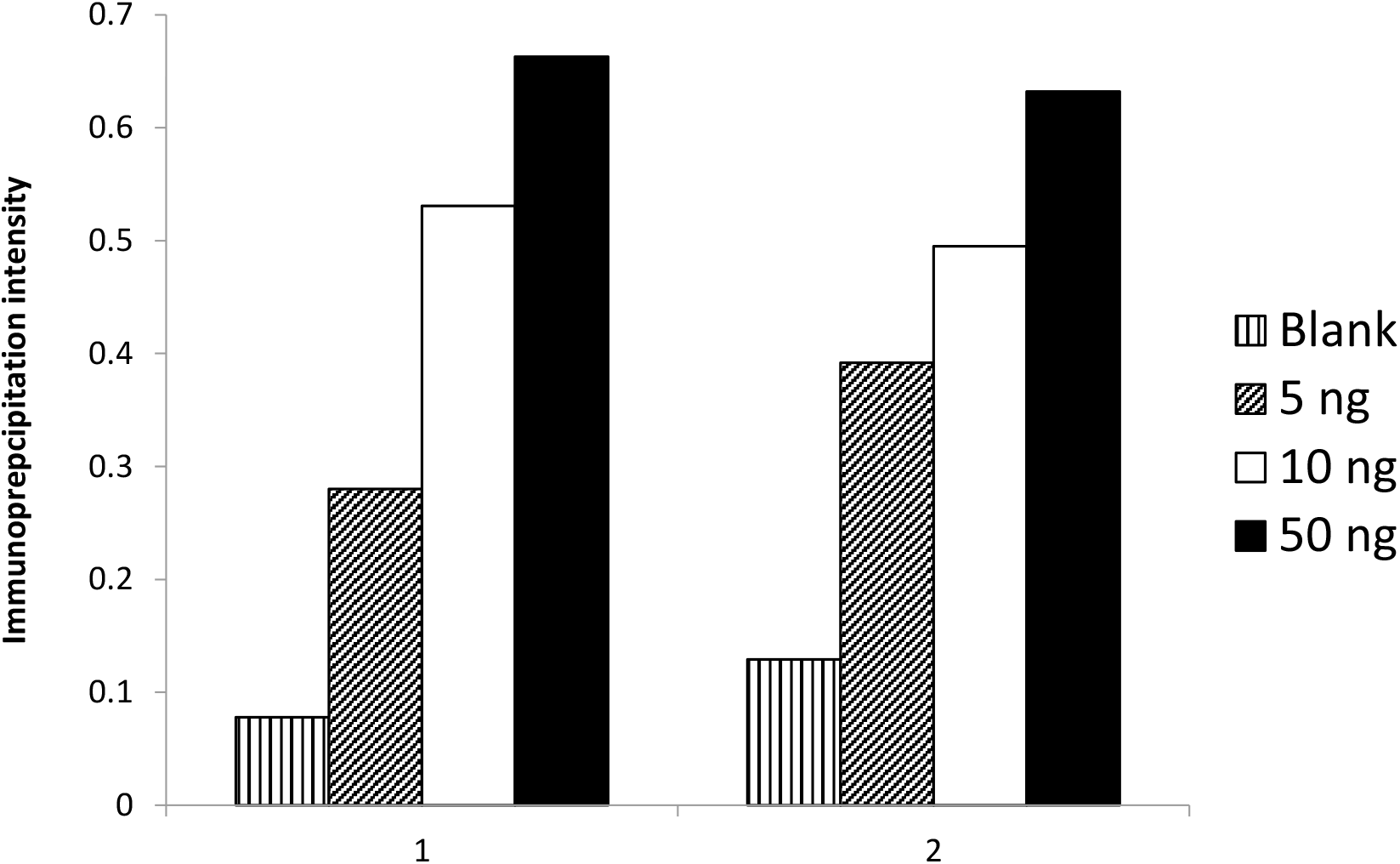
Immunoprecipitation of the SARS-CoV-2 protein containing the S1/S2 boundary furin site by fbAB. A: SARS-CoV-2 protein containing the S1/S2 boundary furin site; B: Peptide containing the SARS-CoV-2 specific furin motif.

Next, fbAB blockage of SARS-CoV-2 furin motif cleavage via enzymatic activity was examined. Dual His- and biotin-tagged peptide containing the SARS-CoV-2-specific furin motif was bound to Ni-coated microplate wells. The wells were then incubated with fbAB, followed by exposure to either of three different enzymes: furin, trypsin, or human nasal swab sample. Cleavage at the furin motif will remove the C-terminal portion of the peptide, resulting in inhibition of streptavidin-HRP binding to biotin and reduced signal intensity. As shown in Figure 6, fbAB effectively blocked the cleavage of the SARS-CoV-2 furin motif by purified furin enzyme at various doses (2-4 U). Peptide pre-incubated with fbAB prior to peptide binding to the assay wells was similarly afforded protection against cleavage by furin (data not shown). By comparison, SARS-CoV-2 neutralization antibody (SnAB), which was used as a control and targets the spike RBD, showed minimal blocking of furin-mediated cleavage. fbAB also downregulated cleavage mediated by the serine protease trypsin (Figure 7), although to a lesser extent when compared with furin. PCs from human nasal swab samples displayed a high cleavage percentage (∼80%) of the furin motif, which was decreased by approximately 30% in the presence of fbAB (Figure 8).

**Figure 6.**
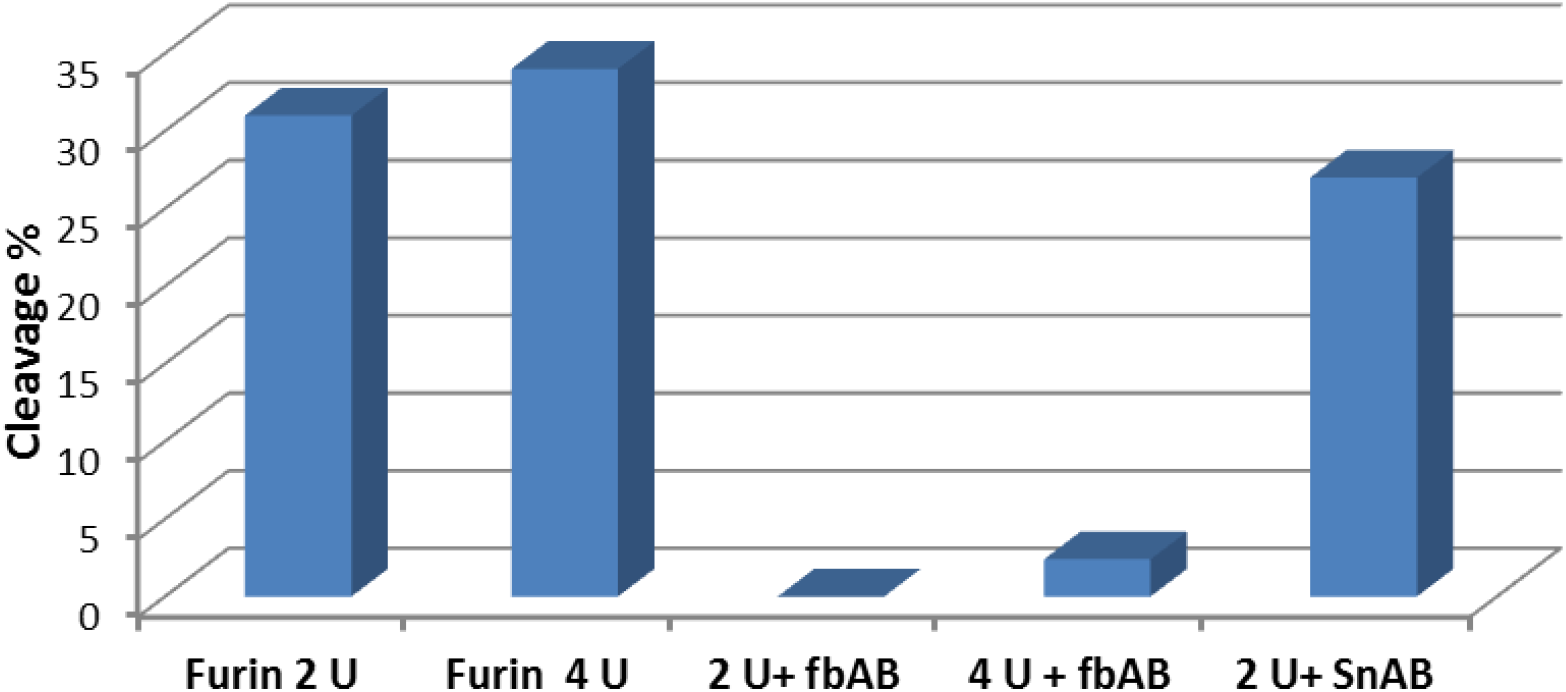
fbAB blockage of SARS-CoV-2 furin motif cleavage by furin. Furin concentration = 2-4 U/well. fbAB and spike protein neutralization antibody (SnAB) concentration = 200 ng/well.

**Figure 7.**
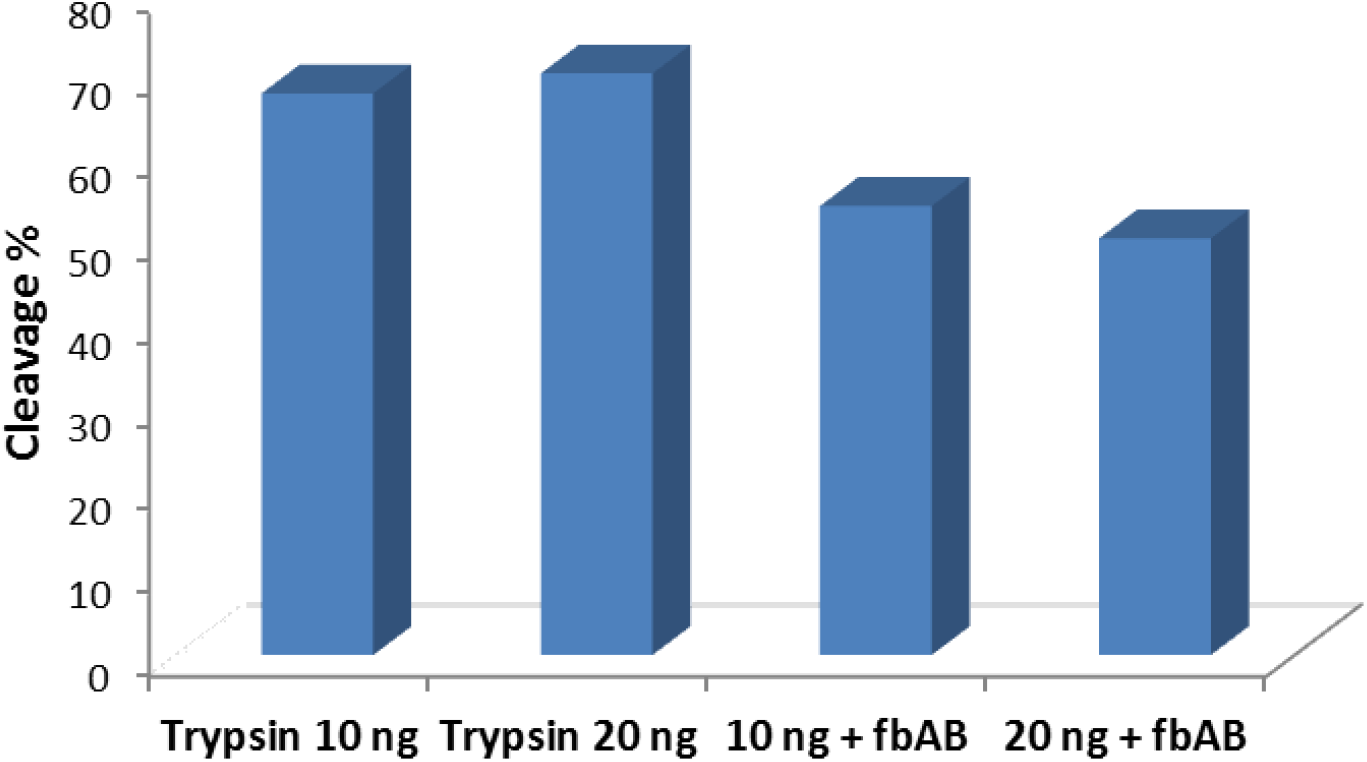
fbAB blockage of SARS-CoV-2 furin motif cleavage by facilitating protease trypsin. Trypsin concentration = 10-20 ng/well. fbAB concentration = 200 ng/well.

**Figure 8.**
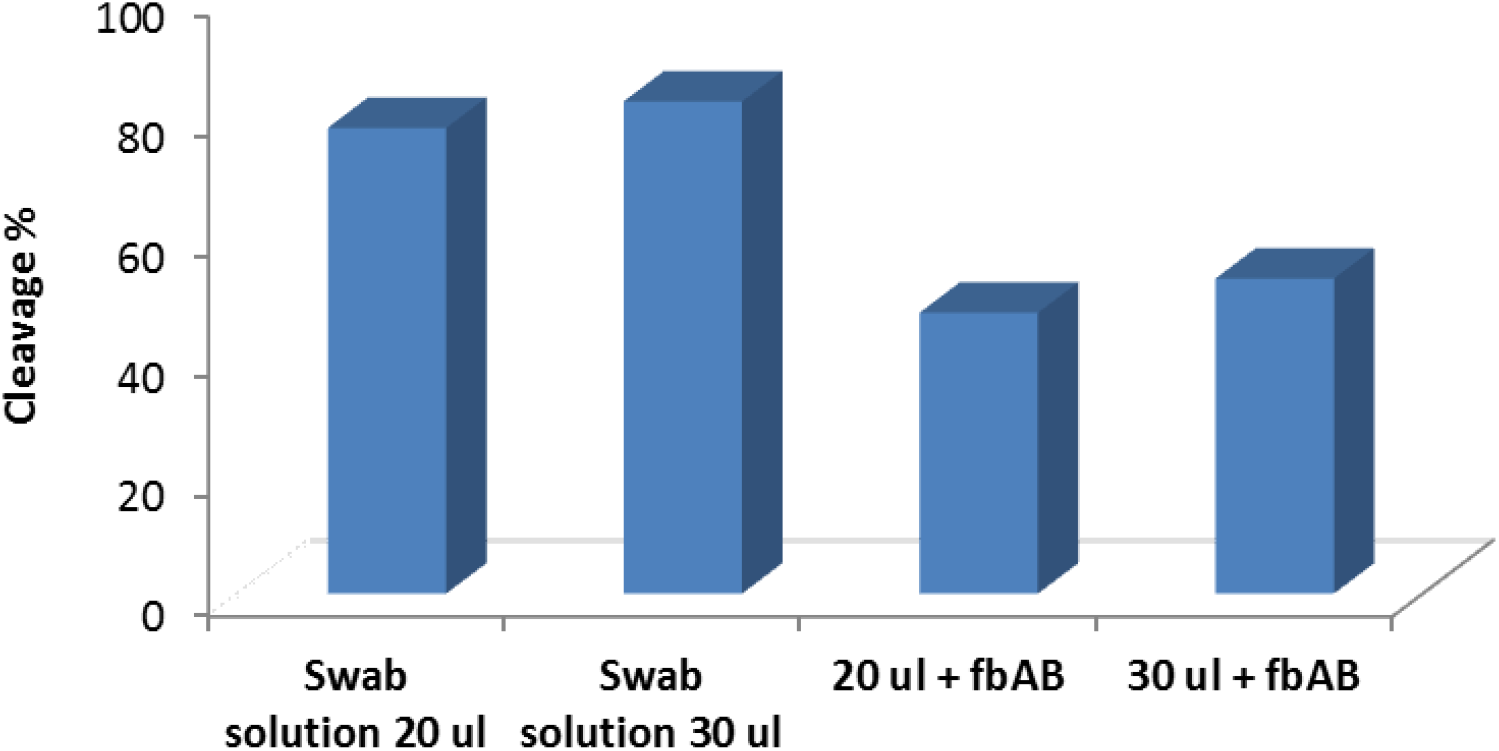
fbAB blockage of SARS-CoV-2 furin motif cleavage by human nasal swab sample. Human nasal swab sample was released into 300 µl of furin assay buffer and 20-30 µl of sample solution was used for the assay. fbAB concentration = 200 ng/well.

As cleavage of the S protein at the S1/S2 site is required for SARS-CoV-2 binding to the host ACE2 receptor for cell entry, we tested the effect of fbAB on spike-ACE2 interaction in SARS-CoV-2 spike protein-coated wells. As shown in Figure 9, fbAB blocked the binding of ACE2 to S protein in a dose-dependent manner, with >60% of ACE2 binding activity diminished at an fbAB dose of 40 nM and almost complete inhibition of spike-ACE2 binding at 80 nM of antibody.

**Figure 9.**
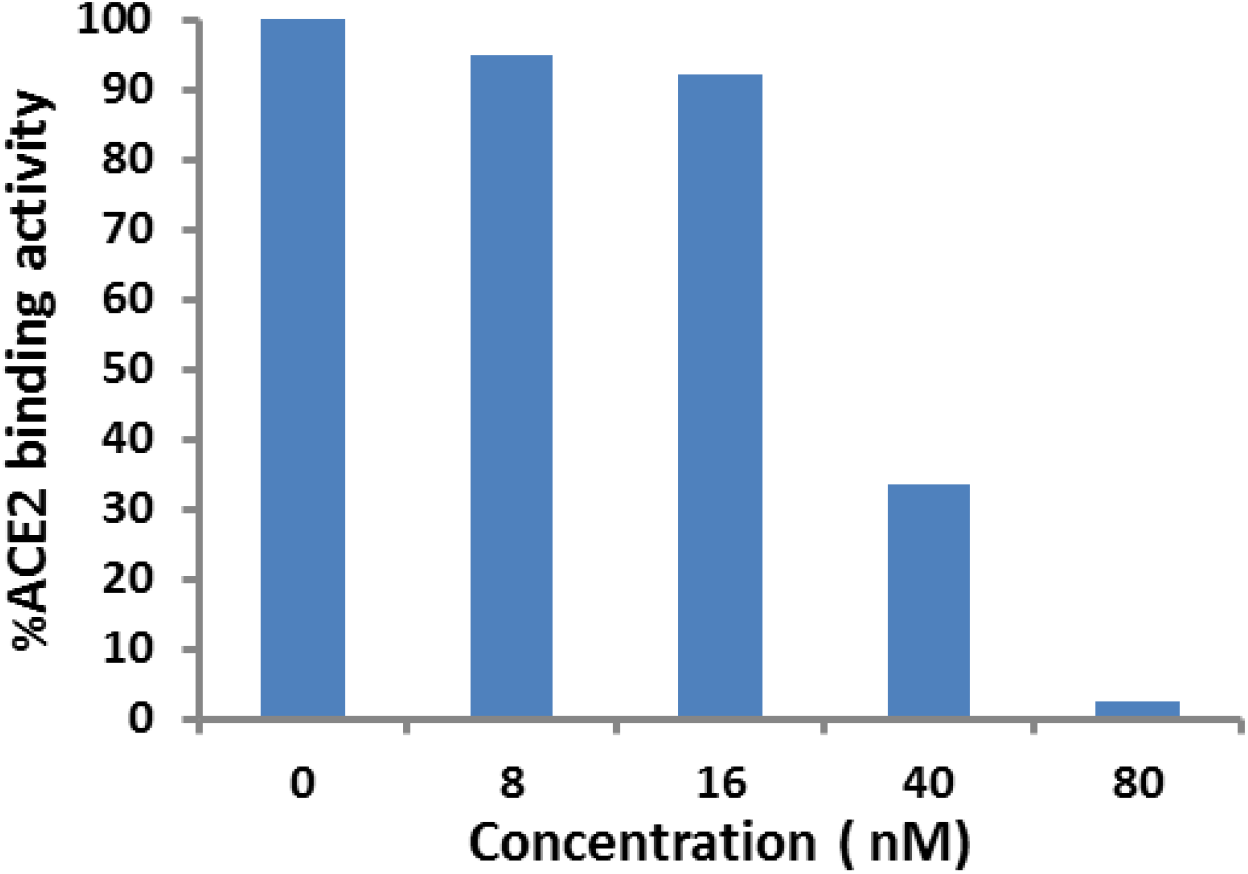
Reduction of ACE2 binding to SARS-CoV-2 spike protein by fbAB blocking the SARS-CoV-2 furin motif at different concentrations. Coated SARS-CoV-2 spike protein concentration = 50 ng/well. ACE2 concentration = 100 ng/well.

## Discussion

During infection, the cell entry mechanism of SARS-CoV-2 involves direct contact with the host ACE2 receptor, facilitated by the RBD within the S1 subunit of the S protein, and proteolytic cleavage of the S1/S2 multibasic cleavage site by the cell surface transmembrane protease serine 2 (TMPRSS2) and cellular cathepsin L [**2**,**3**,**4**]. The S1/S2 junction also harbors a unique furin cleavage site that can be cleaved by furin and other PCs in lieu of TMPRSS2/cathepsin L activity, which may enhance infectivity [**1**,**5**]. A recent study showed that a furin cleavage site-deleted SARS-CoV-2 mutant exhibited reduced replication in human respiratory cells and attenuation of viral pathogenesis in in vivo models [**6**]. The widespread expression of furin, especially in the human lung [**7**,**8**], makes the furin cleavage site an ideal target for therapeutic intervention.

Through use of a synthesized peptide containing the SARS-CoV-specific furin cleavage sequence (entire 20 amino acid motif, Figure 1) as an antigen, a high titer antibody was generated with high sensitivity and specificity for the S1/S2 boundary furin site. This furin site blocking antibody (fbAB) was further examined using a series of innovative ELISA-based colorimetric assays to rapidly and reliably measure its efficiency at blocking cleavage and receptor interaction. fbAB demonstrated effective blockage of SARS-CoV-2 spike protein cleavage caused by purified furin enzyme, the serine protease trypsin, and human nasal swab specimens. It was shown that in both asymptomatic and symptomatic COVID-19 patients, nasal samples have yielded higher viral loads than throat samples [**9**], indicating the nasal epithelium as a portal for initial infection and transmission and as a dominant location for viral replication through pre-activation by PCs from both host and bacteria in the nasal cavity. Based on our results, human nasal swab specimens yielded 80% cleavage of the assay substrate containing the SARS-CoV-2-specific furin motif, confirming this sample type as a valid biological source of abundant PC activity.

It should be noted that in addition to blocking furin-mediated cleavage of bound target, fbAB pre-incubated with peptide containing the SARS-CoV-2-specific furin motif, before immobilization on the well surface, was also effective at inhibiting enzyme cleavage. As virus particles are mobile targets within the systemic circulation, it was necessary to confirm that antibody interaction with unbound target in suspension status could reduce formation of cleaved product as seen with immobile peptide.

Given the importance of the spike/furin/ACE2 signal axis in the infection pathway of SARS-CoV-2, the ability of fbAB to disrupt binding interaction between S protein and its host receptor is critical to attenuating the spread of disease. fbAB demonstrated very high potency in blocking the binding of ACE2 to S protein-coated microplate wells, effectively reducing the percentage of receptor binding activity to a nearly undetectable level at a submicromolar antibody concentration.

Current approaches to blocking or reducing furin site cleavage of target proteins are predicated on the direct inhibition of furin or furin-like enzymes. Such inhibition is generally achieved by naturally-occurring macromolecular protein-based inhibitors (e.g., serpin A1-antitrypsin) or small molecule chemical inhibitors (e.g., pure peptide, peptide mimetics, and nonpeptidic compounds). As these inhibitors are selective for the proteases themselves rather than specific sites of proteolytic activity, their inhibitory effect is limited with regard to preventing furin site cleavage exclusively at a precise location such as the S protein of SARS-CoV-2. Furthermore, furin and furin-related PCs are widely distributed in various human tissues. The inhibition of host proteases could non-specifically damage the normal functionality of proteins that require activation by these enzymes. Therefore, our strategy of implementing an antibody to expressly block the SARS-CoV-2-specific furin site from cleavage by furin and facilitating proteases would be a selective option in controlling the activation of SARS-CoV-2 spike protein and the spread of the virus.

## Notes

### Competing Interest Statement

The authors have declared no competing interest.

